# Scalable volumetric imaging for ultrahigh-speed brain mapping at synaptic resolution

**DOI:** 10.1101/240770

**Authors:** Hao Wang, Qingyuan Zhu, Lufeng Ding, Yan Shen, Chao-Yu Yang, Fang Xu, Chang Shu, Yujie Guo, Zhiwei Xiong, Qinghong Shan, Fan Jia, Peng Su, Qian-Ru Yang, Bing Li, Xiaobin He, Xi Chen, Feng Wu, Jiang-Ning Zhou, Fuqiang Xu, Hua Han, Pak-Ming Lau, Guo-Qiang Bi

**Affiliations:** Hefei National Laboratory for Physical Sciences at the Microscale, and School of Life Sciences, University of Science and Technology of China, Hefei, Anhui 230027, China.; CAS Key Laboratory of Brain Function and Disease, and School of Life Sciences, University of Science and Technology of China, Hefei, Anhui 230027, China.; Institute of Automation, Chinese Academy of Sciences, Beijing 100190, China.; University of Chinese Academy of Sciences, Beijing 100049, China.; School of Information Science and Technology, University of Science and Technology of China, Hefei, Anhui 230027, China.; National Engineering Laboratory for Brain-inspired Intelligence Technology and Application, University of Science and Technology of China, Hefei, Anhui 230027, China.; Wuhan Institute of Physics and Mathematics, Chinese Academy of Sciences, Wuhan 430071, China.; CAS Center for Excellence in Brain Science and Intelligence Technology, Shanghai, 200031, China.

## Abstract

We describe a new light-sheet microscopy method for fast, large-scale volumetric imaging. Combining synchronized scanning illumination and oblique imaging over cleared, thick tissue sections in smooth motion, our approach achieves high-speed 3D image acquisition of an entire mouse brain within 2 hours, at a resolution capable of resolving synaptic spines. It is compatible with immunofluorescence labeling, enabling flexible cell-type specific brain mapping, and is readily scalable for large biological samples such as primate brain.

---

Large scale 3D imaging has become an increasingly important approach in biology, especially for the study of brain circuits ^1^. Systematic brain mapping efforts are building the foundations for our understanding of brain architecture ^2-4^. Meanwhile, targeted labeling and brain-wide tracing together with specific manipulations are revealing circuitry mechanisms underlying brain functions ^5-8^. The rapid expansion in both fronts demands efficient imaging methods. Traditionally, 3D information of fixed brains and other biological samples was obtained by reconstructing series of 2D images of mechanically cut thin sections. Using such samples with fluorescent protein expression or immunofluorescence staining, epifluorescence or confocal microscopy can resolve fine structures such as neuronal synapses ^9^. However, the low throughput of this approach makes whole-brain data reconstruction and analysis a tedious task. Several techniques with automated serial sectioning and imaging methods were developed to overcome this problem ^9-15^. Recent advances along this line was able to push the imaging speed to ~3 days per mouse brain at synaptic resolution ^14^, or ~2.4 hours per mouse brain but at a reduced resolution ^15^. Still, it is highly desirable to be able to obtain high resolution images at very high speed for high volume tasks such as whole brain mapping of many animals or of large primates.

The past decade has also seen various tissue clearing techniques that greatly improve the transparency of brain samples ^16^. Combined with these techniques, serial light-sheet microscopy enables fast but low resolution volumetric imaging of cleared entire brains without mechanical sectioning ^8,17,18^. At the cellular resolution, such a method can achieve imaging of the whole mouse brain within 2 hours ^8^. Light-field microscopy can also be used for fast imaging of cleared brains at cellular resolution ^19^. Importantly, several clearing methods are compatible with immunofluorescence staining, allowing for imaging structures not labeled by genetically encoded fluorescent proteins ^20,21^, even though such staining usually requires prolonged time to complete because of limited diffusion rate of antibodies.

Here, we report a new, scalable strategy implementing a “volumetric imaging with synchronized on-the-fly-oblique-scan and readout” (VISoR) technique for high speed three dimensional imaging of large samples at high resolution. For small samples such as cultured cells ^22^ and *Drosophila* larvae ^23^, state-of-the-art light-sheet microscopy methods have already achieved high-resolution and camera-limited imaging speed. However, these methods cannot be used for larger samples because they rely on fast movement of imaging section across the sample along the axial direction; the range of such movement is thus limited by the working distance of the imaging objective. Our solution is to move the sample horizontally under the V-shape objectives over nearly unlimited range, while using single-pass beam scan to eliminate motion blur, thus allowing for camera-limited high-speed imaging of large samples without compromising image quality. As a proof of principle, we designed and built a system implementing VISoR that can complete image acquisition of an entire mouse brain within 2 hours at 0.49×0.49×3.5 μm^3^ voxel resolution, with individual synaptic spines in cortical neurons readily visible. We have also developed an optimized pipeline for thick brain slice sample preparation as well as programs for semi-automated 3D reconstruction and analysis. The procedure is readily compatible with antibody staining that can be completed within a few days, thus enabling versatile high-throughput cell-type specific 3D brain mapping.

The basic configuration of our system consists of an illumination objective through which a scanning light-beam enters a thick slice sample at a 45-degree angle to the sample surface, and an imaging objective positioned perpendicular to the illumination plane (Fig. 1a,b), similar to the V-shaped light-sheet microscope typically used for cellular imaging ^24^. Both objectives are immersed in refractive index-matched solution (Methods and Materials) so that high quality images of oblique samples sections can be formed at the camera sensor (Supplementary Fig. 1). The geometry and working distance of the objectives constrain the imageable sample thickness to ~300 μm (Fig. 1b). The light path is configured such that the effective imaging area on the object plane (corresponding to half of the camera sensor, ~1000 X 2000 pixels) covers an oblique section of ~420 μm X 1000 μm using a 20X imaging objective with system magnification reduced to ~13X (Fig. 1c).

**Figure 1.**
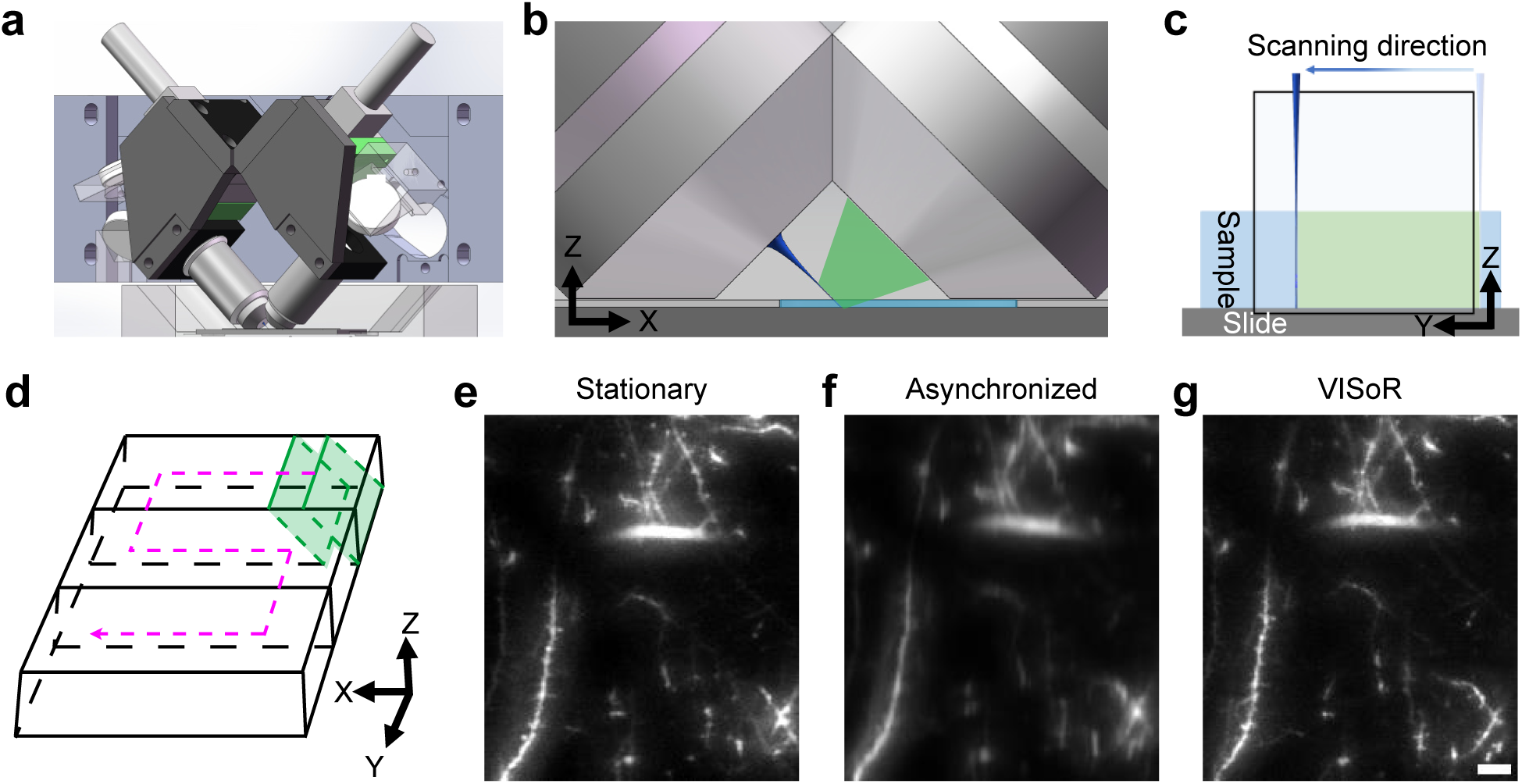
The mechanism and implementation of VISoR. **(a)** The core optics of the system consists of two perpendicular objective lenses mounted 45 degrees oblique to the sample surface. **(b)** Enlarged view of the working area, showing the geometric configuration of the two objectives over the sample slice (light blue), along with the illumination beam (dark blue) and fluorescence collection path (green). **(c)** The imageable area in the object plane (in conjugation with the camera sensor) is illuminated by the scanning light beam synchronized to camera readout. Note that only half of the sensor area is utilized for acquisition of single-channel images. **(d)** Volumetric imaging is implemented by synchronous single-pass beam-scan and on-the-fly readout over the surface of a sample slice with a zigzagged motion path. **(e-g)** Fluorescence images of the same sample area obtained with stationary imaging **(e)**, non-synchronized on-the-fly imaging in “light-sheet” mode **(f)**, and VISoR imaging **(g)**. Scale bar: 10 μm.

To achieve fast volumetric imaging over a large range, we configured the opto-mechanical control of the system to allow horizontal non-stop smooth motion of the sample stage with on-the-fly image readout of oblique optical sections continuously (Fig. 1c, d). This approach has also been used in the recent “open-top light-sheet microscopy” and “oblique light-sheet tomography”. ^25,26^. However, because the movement is not along the axial direction, the lateral component motion naturally cause smear (motion blur) of the acquired images that is proportional to the speed of moving stage (Fig. 1e,f). To overcome this problem, we implemented a thin excitation beam (3-5 μm diameter) to scan through the continuously moving sample, synchronized with image acquisition cycle. At 100 Hz scanning and imaging cycle with the stage moving at 0.5 mm/s, the effective excitation time for each voxel is less than 100 μs, within which the motion is less than 0.05 μm. With this strategy, we eliminate virtually all motion blur and achieve high image quality indistinguishable from stationary imaging, with dendritic spines of cortical neurons clearly visible (Fig. 1e-g). In principle, motion blur can also be reduced using strobe light illumination in a regular light-sheet configuration. This however requires much higher light power to achieve similar illumination. The synchronized beam scan is thus also optimized in light efficiency.

Our prototype system allows for up to 200 Hz scanning-imaging cycle at 1 mm/s translation for single channel, or 100 Hz at 0.5mm/s for dual-channel imaging with a single camera and a channel splitter. The system can run mostly continuously at the highest possible data rate of the camera, up to 400 million pixels per second, equivalent to ~0.3×1.0×1.0 mm^3^ volume imaged per second at ~0.49×0.49×3.5 μm^3^ voxel resolution. To test the capacity of this new system in whole-brain imaging, we started with fixed brains from *Thy1*-YFP-H mice ^27^. An embedded mouse brain was cut into ~50 slices of 300 μm thickness each, to match the imaging depth of the optical system (Fig. 1b). A pipeline was developed to optimize brain fixation, slice cutting and handling, as well as tissue clearing and mounting procedures (Supplementary Fig. 2a). We also developed a system to automatically detect the edges of all mounted brain slices, so that the imaging system can directly move to the target based on correlated coordinates. (Supplementary Fig. 2b). Because of the oblique configuration of excitation and imaging light paths relative to the sample surface, tissue clearing and refractive index matching are critical for high-quality imaging, as mismatched immersion solution could lead to severe aberration (Supplementary Fig. 3). With optimized sample preparation and imaging conditions, high quality volumetric image series or “columns” can be obtained (Supplementary Video 1). Adjacent image columns were taken with about 10% overlap, in order to facilitate the alignment and reconstruction of the entire slice (Supplementary Videos 2 and 3). In our experiments with 20x objective (final 13.3x magnification), 100 Hz scanning, and 0.5 mm/s stage speed, corresponding to ~0.49×0.49×3.5 μm^3^ voxel resolution, volumetric data acquisition of a complete set of about 45-50 300-μm slices of a whole mouse brain can be completed in about 2 hours.

Reconstruction of the whole brain volumetric imaging data was performed with custom software that used minimal morphing to compensate for structural deformations during sample preparation (See Methods and Materials). Aside from the interfaces between adjacent slices that exhibit some misalignments, in part due to structural alterations during sample preparation, the reconstructed whole brain shows well-preserved, consistent 3D structural details (Fig. 2a–d, Supplementary Video 4). Neuronal axons and dendrites as well as dendritic spines can be clearly visualized (Fig. 2e–g, Supplementary Fig. 4). Similar results were obtained from all four *Thy1*-YFP-H mice brains tested. In addition, high quality data were also acquired for various brain samples labeled using neurotropic viruses, including a Semliki Forest virus (SFV) variant that sparsely labels neurons in the hippocampus (Supplementary Fig. 5), and a vesicular stomatitis virus (VSV) that labels dopaminergic neurons in the ventral tegmental area (VTA) through nerve terminal infection at the nucleus accumbens (NAc) (Supplementary Fig. 6, Video 5).

**Figure 2.**
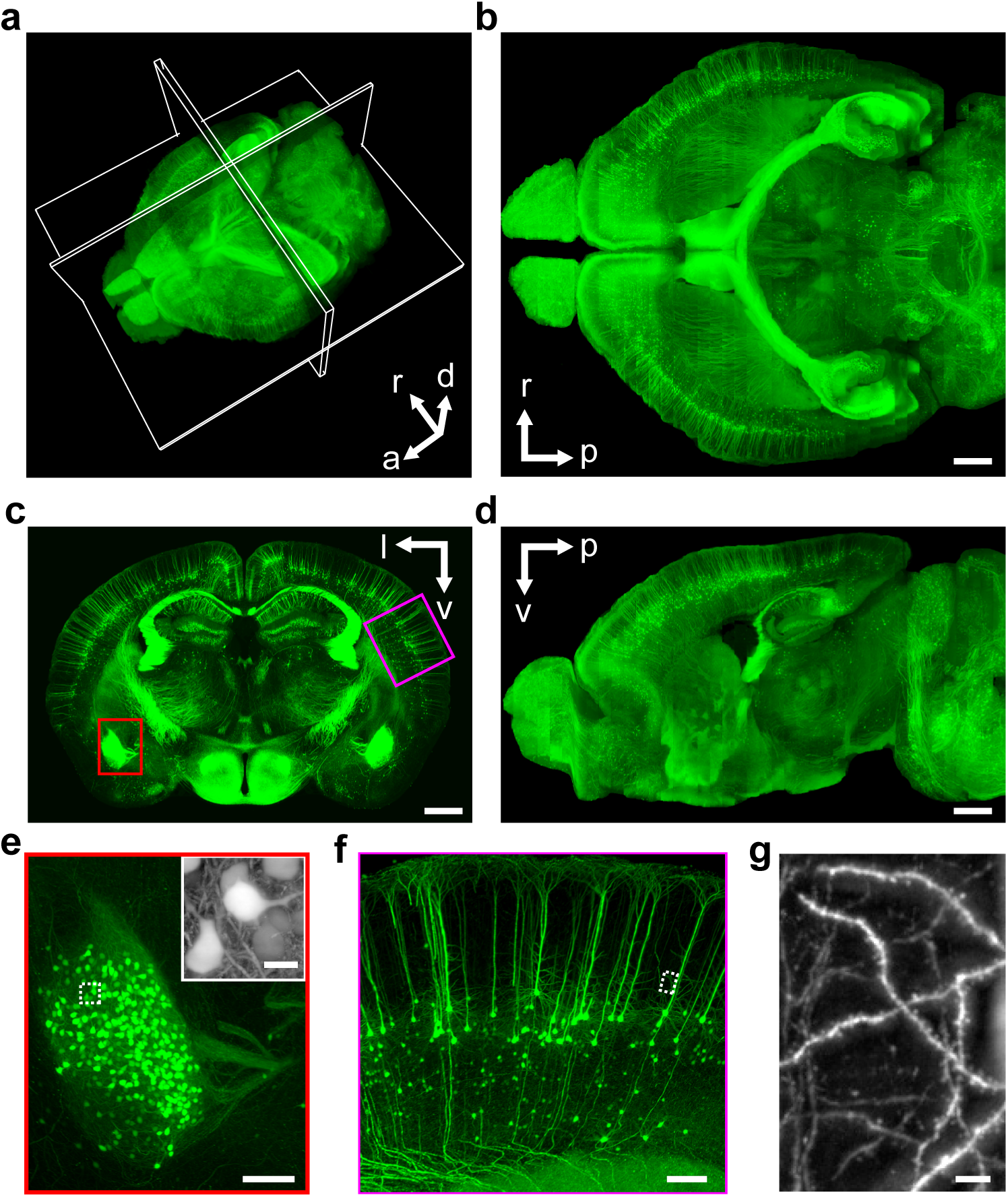
Whole brain VISoR imaging of a Thy1-YFP transgenic mouse. **(a)** 3D reconstructed mouse brain. Maximum intensity projections (MIPs) of a 800-μm horizontal section, a 300-μm coronal section and a 150-μm sagittal sections are displayed in **(b)**, **(c)** and **(d)**, respectively. The position of each section is illustrated in **(a)**. Abbreviations: l, left; r, right; d, dorsal; v, ventral; a, anterior; p, posterior. **(e)** Enlarged view of the red box in **(c)** showing the left amygdala. Inset: an enlarged view of the white rectangle area. **(f)** Enlarged view of the magenta box in **(c)** showing part of the cortex. **(g)** Enlarged view of the white rectangle area in **(f)**. Note that the inset of **(e)**, and **(g)** are MIPs of the corresponding areas with projection direction perpendicular to the imaging plane, i.e. 45° oblique to the slice surface, thus appearing to be different from the direct magnification of the corresponding rectangle areas. Scale bars: **(a)**, **(b)**, **(c)**, **(d)**, 1000 μm; **(e)**, **(f)**, 200 μm; inset of **(e)**, 20 μm; **(g)**, 10 μm.

An advantage of hydrated sample preparation is its compatibility with immunofluorescent staining. This allows for convenient labeling of different cell types or states, enabling flexible cell-type specific brain connectivity and activity mapping. As an example, we used c-fos antibodies to label activated neurons in the brains of CRH-ires-Cre^+/+^::Ai14^+/−^ transgenic mice with or without brief footshock stimuli (Methods and Materials), and performed dual color VISoR imaging and cell counting in different brain areas (Fig. 3). In the paraventricular nucleus (PVN) of hypothalamus where CRH neurons were observed to form a dense cluster (Fig. 3a,b), more than half of these neurons (430 out of 834) in the left PVN of the stimulated animal were c-fos positive, whereas very few cells (37 in total) in the left PVN of the control animal were activated, among them most were CRH neurons (35 out of 771 CRH neurons) (Fig. 3b,c, Supplementary Video 6). This is consistent with the role of CRH neurons of the PVN in stress response ^28^. Interestingly, in the medial amygdala (MeA) where CRH neurons were more dispersed (Fig. 3d), the CRH-expression and c-fos labeling exhibited a near orthogonal pattern in the stimulated animal, with 104 CRH neurons being activated out of 1018 total CRH neurons and 569 total activated neurons (Fig. 3d,e, Supplementary Video 7). Again, few neurons (63 in total) in this area were found activated in the control animal (Fig. 3f). Although more systematic experiments and analyses are needed to understand the activation patterns of CRH neurons in different areas, these results do suggest that they may function very differently in the animal’s stress response. They also demonstrate the effectiveness of our system in combining fast 3D fluorescence imaging with antibody staining to investigate the active roles of specific cell types in brain function.

**Figure 3.**
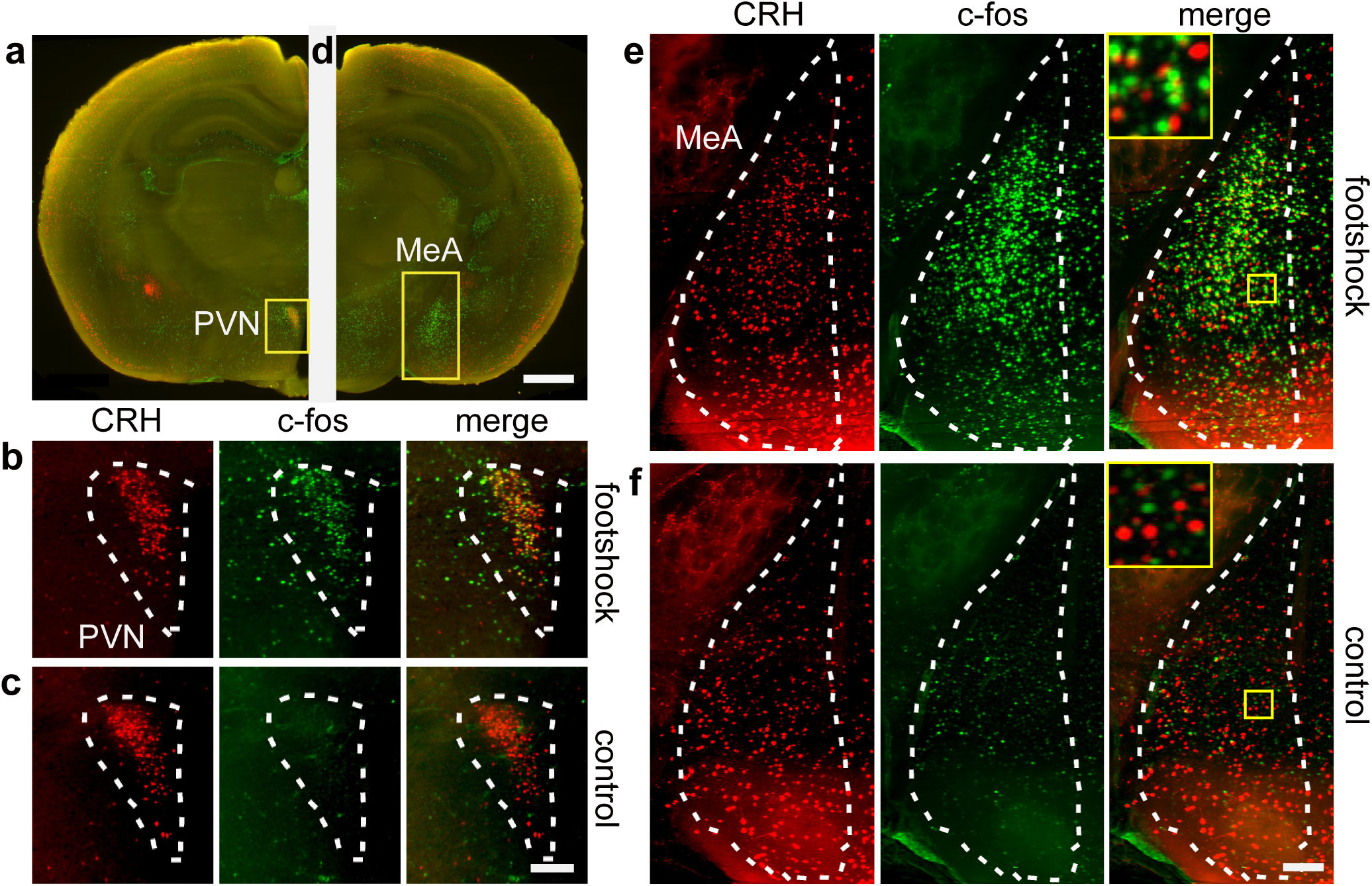
Dual-channel VISoR imaging of c-fos expression revealing differential stress-elicited activation of CRH neurons in different brain areas. **(a)** CRH and c-fos expression in a virtual coronal section (A/P: bregma −0.72~−0.9) of a CRH-ires-Cre^+/+^::Ai14^+/−^ transgenic mouse exposed to footshock stimulation. Green, c-fos immunostaining; red, tdTomato expressed in CRH neurons. **(b)** Magnified view of the PVN area as shown in **(a)** from the same animal. **(c)** Magnified view of the PVN area a control animal not exposed to footshock stimulation. **(d)** CRH and c-fos expression in a virtual coronal section (A/P: bregma −1.58~−1.78) contain the MeA from the same animal as in **(a)**. **(e)** Magnified view of the MeA area as shown in **(d)**. **(f)** Magnified view of the MeA area from a control animal. MIP of thinner (7 μm) virtual sections are used for **(b)** and **(c)** in order to visualize single cells. Scale bars: **(a)**, **(d)**, 100 μm; **(b)**, **(c)**, **(e)**, **(f)**, 100 μm.

With continuous on-the-fly imaging, the speed of the VISoR system is mainly constrained by data acquisition bandwidth. For certain brain mapping applications where synaptic resolution is not critical, a VISoR system can achieve much higher throughput, e.g. one entire mouse brain within 15 minutes at a reduced pixel resolution of ~2 μm. Additionally, the axial resolution effectively limited by beam width may be improved with other illumination strategies such as 2-photon Bessel beam ^29^, or with the dual-view configuration as in diSPIM for cellular imaging ^22^. It is worth noting that the VISoR technique can be implemented in different ways and can find applications in the study of 3D structures of other biological and pathological specimen beyond brain mapping. Meanwhile, the strategy of clearing and staining performed after sectioning considerably reduces sample preparation time, from weeks required for antibodies staining of an intact mouse brain ^20,21^, to a couple of days for staining 300-μm slices. Thus, our approach is ideally suited for whole brain mapping of neuronal connectivity and activity in different levels of details, and is readily scalable to larger samples including primate brains.

## METHODS AND MATERIALS

### Optical setup

The microscope system (Supplementary Fig. 1) consists of a light source module, a scanning light-beam illumination module, an imaging module, a sample chamber for imaging and a 3-axis automatic stage for sample positioning. The light source module combines four compact lasers with wavelength 405nm, 488nm, 561nm and 647nm (Coherent Inc.) and delivers light to the illumination module, which consists of a galvanometer scanning mirror (GVS011, Thorlabs) and relay lens sets. The optical axis of illumination path and imaging path are perpendicular, and 45-degree off the sample surface. (Supplementary figure 1). The illumination and imaging modules have identical front ends, which consist of an objective (UMPLFLN20XW, NA0.5, 20X water immersion, Olympus) and a micropositioner, which allows potential configuration expansion of dual-excitation/dual-imaging. The focal length of the tube lens in the imaging module is 120 mm which provides an effective magnification of 13.3X. A dual-channel splitter (Optosplit II, Cairn Research, UK) is mounted between the imaging module and the sCMOS camera (ORCA Flash 4.0 v2, Hamamatsu) to provide two imaging channels at the same time. A combination of emission filters (Semrock Brightline 452/45, 520/28, 593/46 and Chroma ET705/72 M) are mounted in the Optosplit II. The imaging chamber carries the sample and the refractive index matched immersion solution, and is mounted on the top of a 3-axis motorized stage, which consists of two one-dimensional translation stages (MTS50, Thorlabs) for X/Y-translation and a heavy duty stage for Z-translation.

### Computers, electronics and control programs

All electronic components of the system including lasers, galvanometer scanner, DAQ board (PCI-6722, National Instruments) and camera are controlled by a workstation (dual Xeon CPUs, 64GB RAM, Dell) equipped with 8 terabyte SSD RAID0 array and 10 Gigabit optical fiber networks connecting to large volume storage servers (200 TB, Dell) and computation servers (4 nodes, 28 cores and 128 GB RAM for each node). The whole system is controlled by custom C++ code based on MicroManager ^30^ core. Imaging data is directly acquired into the SSD array, then transferred to the large volume storage server for further reconstruction and quantitative analysis.

### Virus preparation

The viral-like particles of Semliki forest virus SFV-EGFP were prepared as follows. Briefly, SFV-EGFP replicon was modified based on the SFV replicon followed by *in vitro* transcription. The SFV-EGFP replicon RNA and helper RNA were co-transfected into BHK21 cells to package SFV-EGFP. The detail procedures for RNA transcription and virus package were reported in previous studies ^31,32^. The SFV replicon and helper plasmids were kindly donated by Dr. Markus U. Ehrengruber, Department of Biology, Kantonsschule Hohe Promenade, Zurich, Switzerland. The mutated vesicular stomatitis virus rVSV-EGFP-NR7A were prepared as described previously ^33,34^.

### Animal experiments

All animal experiments were carried out following protocols approved by the Institutional Animal Care and Use Committees of the University of Science and Technology of China (USTC) and Wuhan Institute of Physics and Mathematics, the Chinese Academy of Sciences. All mice used in this study were group-housed with 12 hours light/dark cycle (light on at 7 a.m.) with free access of food and water. Eight-week-old male Thy1-YFP-H (Jax: 003782) mice were used for whole brain structural imaging. A male C57BL/6 mouse of the same age was used for SFV injection. A DAT-ires-Cre ^35^ (Jax: 000660) male mouse (8 weeks old) was used for AAV-DIO-BFP-T2A-TVA and VSV-EGFP-NR7A injection. Male CRH-ires-Cre^+/−^::Ai14^+/−^ mice (used at 12 weeks old) were generated by crossing male CRH-ires-Cre+/+ (*Crh^tm^*^1(^*^cre^*^)^*^Zjh^*) ^36^ (Jax: 012704) and female Ai14^+/−^ (Rosa-CAG-LSL-tdTomato-WPRE::deltaNeo) ^37^ (Jax: 007908) mice.

SFV injection was performed as described in previous study^38^. Briefly, the mouse was anesthetized with chloral hydrate (400 mg/kg) and placed in a stereotaxic apparatus. SFV-EGFP (2.6×10^7^ infectious particles/ml) was injected into the dentate gyrus of the hippocampus. The mouse was sacrificed 24 hours later. For VSV tracing experiment, the DAT-Cre mouse was kept under anesthesia as described above. Injection coordinates (in mm) used were: VTA, A/P −3.20 from bregma, L/M −0.40, D/V −4.30; NAC, A/P 1.5 from bregma, L/M −1.1, D/V −4.6. 50nl of 2×10^8^ ffu/mL rVSV-EGFP-NR7A(A/RG) was injected into VTA 2 weeks after 100nl of 2×10^12^ vg/ml rAAV-DIO-BFP-T2A-TVA was injected into NAc. The mouse was sacrificed 5 days after rVSV-EGFP-NR7A(A/RG) injected. These experiments were conducted in a biosafety level 2 laboratory.

For foot-shock stimulation, a male CRH-IRES-Cre^+/−^::Ai14^+/−^ mouse was given 15 trials of electric shocks (2 seconds, 0.7 mA) randomly spaced in 10 minutes in an isolated behavioral test box with video recording, then rested in home cage for 80 minutes before sacrificing and perfusion.

For brain harvesting, mice were first anesthetized by intraperitioneally injection of 1% pentobarbital sodium (80 mg/kg). Fixation solution containing 4% paraformaldehyde (PFA) with or without 0.5-4% acrylamide hydrogel monomer solution (HMS) in PBS (w/v) was prepared and stored at 4°C before use. Brains were harvested and incubated in the same fixation solution at 4°C for overnight to two days.

### Preparation of brain slice samples

Fixed brains were cut into 300-μm-thick slices using a vibroslicer (Compresstome VF-300, Precisionary Instruments). The brain slices were then transferred into the clearing solution for 12~24 hours at 37°C with gentle shaking. The clearing solution was either 4% SDS solution^20^ or 0.5% PBST. After clearing, the slice samples were washed with 0.1% PBST 3 times before being mounted on quartz slides.

For antibody staining, cleared slices were loaded into 12-well plates, permeated and blocked in blocking solution (5% BSA (w/v), 0.3% TritonX-100 in PBS) for 1 hour and then incubated with primary antibody (Rabbit-anti-cfos, Santa Cruz SC-52, lot: K0112, dilution 1:500) in blocking solution for overnight, followed by washing in 0.3% PBST 3 times for 1 hour each. Secondary antibody (AF488-tagged donkey-anti-rabbit IgG, Jackson ImmunoResearch Labs 711-545-152, lot: 129588, dilution 1:200) was 5 hours in blocking solution, followed by washing in 0.3% PBST. The plates were kept on a gentle bouncer at room temperature for all above steps.

### Fluorescence imaging

Samples on imaging slides were immersed in refractive index matching solution ^39^ (80% Iohexol in PBS, Sigma (D2158)) 4 hours before being mounted in the imaging chamber filled with refractive index matching solution. 16-bit images were acquired with synchronized 100 Hz illumination beam scan and 100 frames per second camera readout, with the sample stage moving at 0.5 mm/s in X direction (Fig. 1d). The resultant voxel size is ~0.49×0.49×3.5 μm^3^.

### Whole brain image reconstruction

Image reconstruction and brain registration were performed using custom programs in ImageJ^40^, MATLAB (2016b, Mathworks Inc.), or C++. The obliquely acquired images were first stacked into columns according to their real world coordinates, then the columns were stitched together to form whole slices based on the coordinates and correlations in their overlay regions.

For the reconstruction of Thy1-YFP brains, we first enhanced contrast of each slice and then reconstructed the whole brain structure in four steps. (1) The top and bottom surfaces of each brain slice were fit into flattened planes by linear regression and interpolation. Structures like ventricles were avoided in the flattening procedure. (2) Edges and textures were detected in the opposing surfaces of adjacent slices, with correspondences between the two surfaces extracted using SIFT-flow ^41^. (3) To limit the error accumulation and propagation through multiple slides, we adjusted the correspondences under two additional constrains over all slides, based on the assumptions of smoothness and small deformation, i.e., the displacements of the neighboring correspondences should be similar and the displacement vector for the correspondences to should as small as possible. (4) With the extracted and adjusted correspondences, moving-Least-Square (MLS) method ^42^ is used to warp each slice with minimal distortion in sizes and shapes of the slices.

### Image visualization

We developed an adaptive tone mapping strategy for efficient visualization of global and local structures in 8-bit display based on the original 16-bit data. The original high-dynamic-range image is decomposed into a base layer and a detail layer through bilateral filtering ^43^. The dynamic range of the base layer is compressed by a global operator, and the result is then recombined with the uncompressed detail layer. To cope with the extremely high local contrast exhibited by the fluorescent protein labeled neurons in the brain image, we propose a progressive tone mapping strategy that compresses the dynamic range of the base layer step-by-step, and the parameters used in bilateral filtering are adaptively determined according to the image content. This allows for capturing the details of both high brightness structures such as neuronal soma or fiber bundles and low brightness structures like dendritic spines or axon terminals in the same field of view.

### Cell detection, counting, and brain registration

Quantitative analyses were performed using custom programs coded in C++ and Python incorporated with OpenCV 3 library. The soma of tdTomato-expressing neurons or the nuclei of c-fos positive cells were detected by finding the local maxima of Difference-of-Gaussian (DoG) in original 2D images, then segmented using watershed algorithm based on local threshold. The 3D somas or nuclei were reconstructed by connecting detected segments of the same cell nucleus from neighboring images. The signals of c-Fos or CRH positive cells are detected and located use a custom C++ code. The cell detecting algorithm consisted of three steps: (1) Finding all local maxima of DoG (Difference of Gaussian) of 4x downsized original image, and dividing them using watershed algorithm. (2) Extracting image patches of surrounding of each local maxima, and find the area of cell by threshold. The side length of patches are 3 times of predefined cell size. The thresholds is set to the average of the value of the local maxima and the mean value of edges of the patch. (3) Aligning areas of cells between neighboring images. After detection, the position, volume, eccentricity and intensity of cells are calculated, then compared to predefined ranges to validate the detected cell.

Automatic brain registration and annotation were performed using a procedure similar to that previously reported ^21^. The 3D volumes were downscaled to 25 × 25 × 25 μm pixel size, then flattened and stitched to reconstruct the whole brain. Elastix (http://elastix.isi.uu.nl) was used to register the reconstructed brain to the reference autofluorescence brain (http://www.idisco.info). The position of cells in whole brain were refined by the transform parameters of brain registration.

## Code availability

Codes for data analysis are available from the authors upon request.

## Data availability statement

Original data are available from the authors upon request.

## ACKNOWLEDGEMENTS

We thank Meiyu Shi, Keming Zhang and Dasheng Bi for their help and suggestions, Dr. Markus Ehrengruber for providing SFV replicon and helper plasmids. This work was supported in part by the Strategic Priority Research Program of the Chinese Academy of Sciences (XDB02050000, XDB02060000, XDB02030000), the National Natural Science Foundation of China (91232722, 91432305), and the National Basic Research Program of China (2013CB835101).

## AUTHOR CONTRIBUTIONS

H.W., Q.Z. and G.-Q.B. invented VISoR, H.W., Q.Z. and L.D. built the system, H.W., Q.Z., P.-M.L. and G.-Q.B. designed the experiments, H.W, L.D., Y.S., C.-Y.Y., Q.-R.Y. and B.L. performed the experiments, H.W, L.D., C.-Y.Y., F.X.(2), C.S., Y.G., Z.X., X.C., F.W. and H.H. designed analysis software and analyzed the data, Q.S. and J.-N.Z. provided brain samples of control and foot-shocked CRH-tdTomato transgenic mice, F.J., P.S., X.H. and F.X.(7) provided brain samples infected with tracing viruses, H.W., F.X.(2), Y.G., P.-M.L. and G.-Q.B. wrote the manuscript with inputs from all authors.

---

**Supplementary Figure 1.**
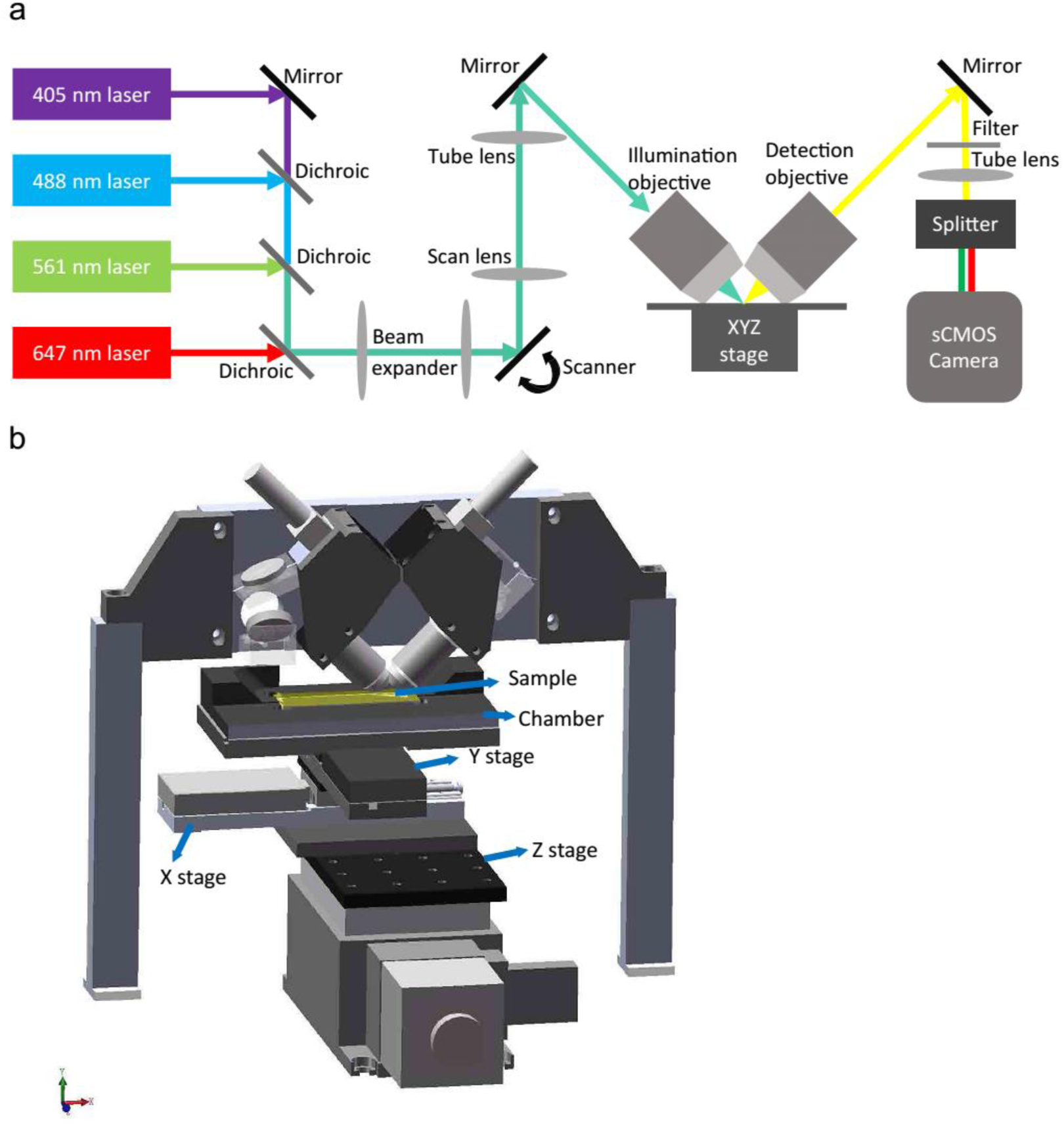
Schematic diagram of the optical system implementing VISoR. **(a)** The illumination and detection objectives are mounted perpendicular to each other and 45 degree to the surface of the sample. The excitation light beam through the galvanometer scanner and illumination objective forms a scanned light sheet inside the sample that matches exactly the imaging focal plane of the detection objective. The sample chamber mounted on a motorized XYZ stage contains imaging solution with refractive index matched to that of the sample. The detection module consists of emission filters, a tube lens, a dual-channel splitter and a sCMOS camera. Camera readout, laser output and galvanometer scanning are synchronized with an NI DAQ board. **(b)** Optical implementation of the VISoR system.

**Supplementary Figure 2.**
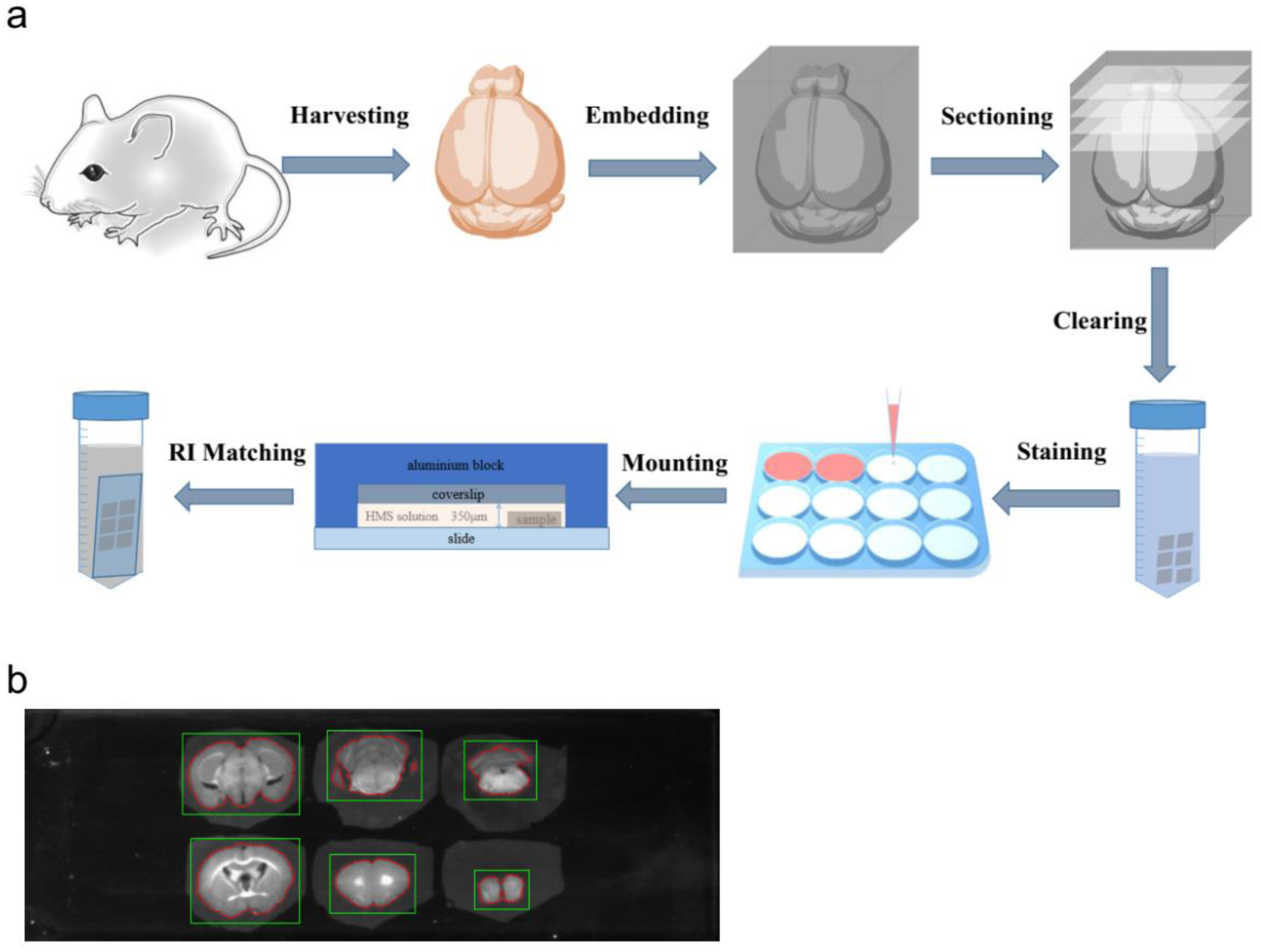
Pipeline for sample preparation. **(a)** The procedure of sample preparation includes following key steps: brain harvesting, embedding, sectioning, clearing, staining (optional), mounting and refractive index matching. With antibody staining, the total time for sample preparation is about 4 days. **(b)** The boundary of each slice (red cycle) is detected by analyzing an pre-acquired image before refractive index matching for further imaging area (green rectangle) determination.

**Supplementary Figure 3.**
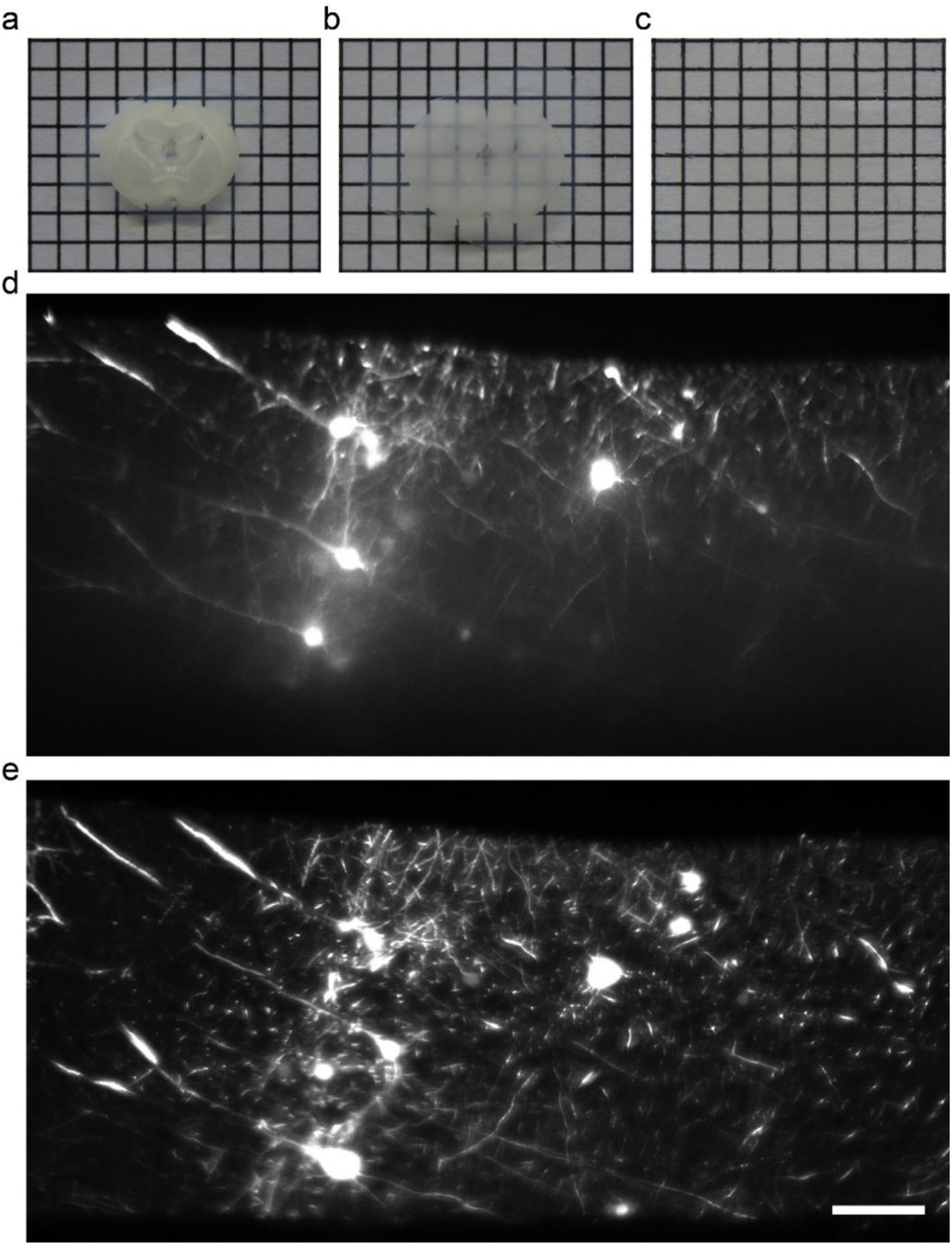
Refractive index matching is essential for high quality imaging. **(a-c)** A 300-μm thick brain slice is mostly opaque in PBS **(a)**, becomes translucent after clearing with 4% SDS for 24 hours **(b)**, and fully transparent after refractive index matching in 80% iohexol solution **(c)**. **(d)** An example VISoR image of a cleared brain slice in PBS shows strong scattering and aberration especially at the lower part of the slice. **(e)** A nearby section of the same sample after refractive index matching, taken under otherwise the same imaging conditions as in **(d)**, showing considerably reduced scattering and aberration. Scale bars: **(a)**, **(b)**, **(c)**, 2 mm per grid; **(d)**, **(e)**, 100 μm.

**Supplementary Figure 4.**
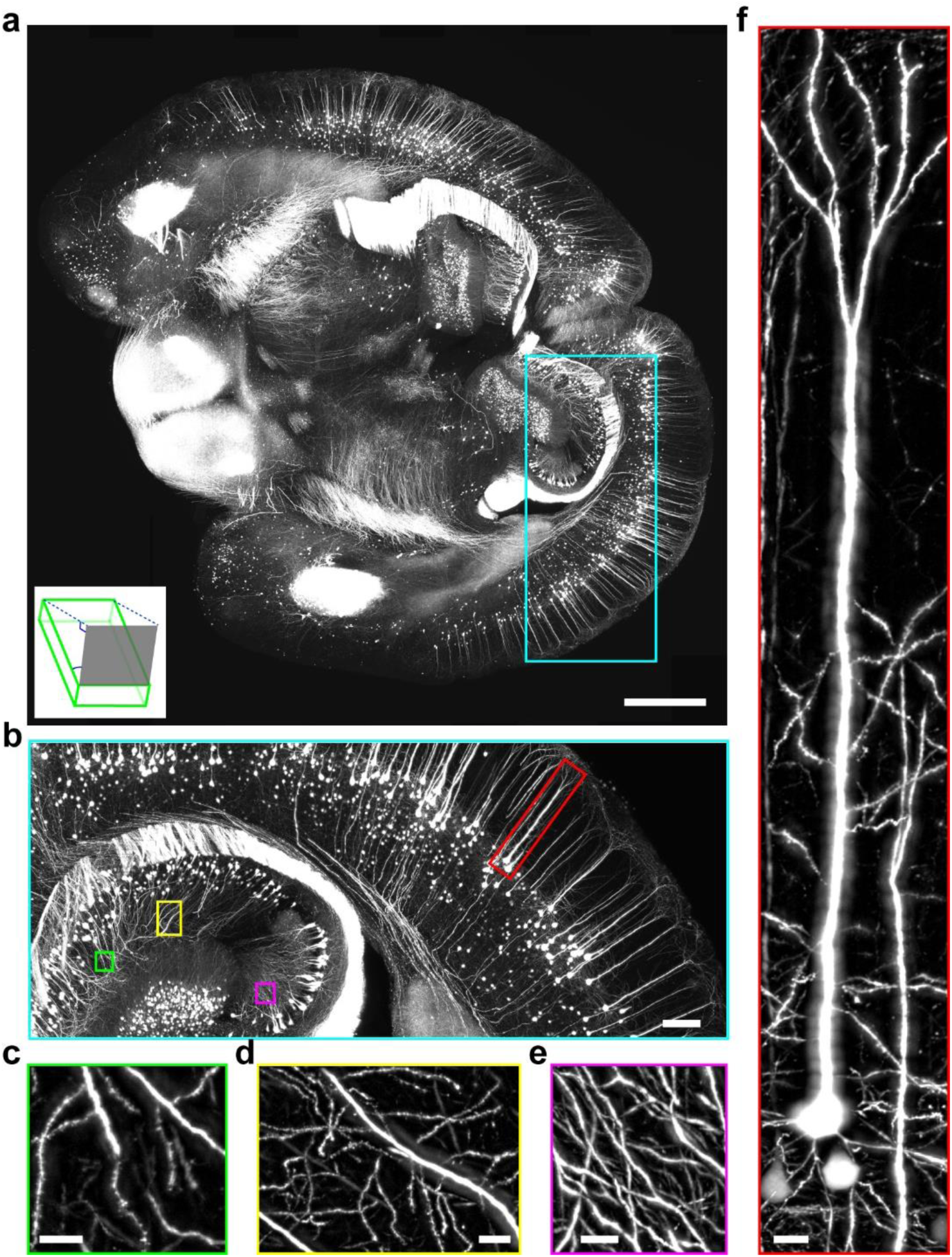
Examples of high-resolution VISoR images. **(a)** A 300-μm-thick slice viewed as Maximum intensity projection (MIP) to the imaging plane (gray plane in the inset), 45° from the slice (green volume in the inset) surface. **(b)** Enlarged view of the cyan rectangle area in **(a)**. **(c-e)** Different hippocampal areas in **(b)** indicated by the green **(c)**, yellow **(d)** and magenta **(e)** rectangles. **(f)** A cortical pyramidal neuron indicated by the red rectangle in **(b)**. Note that in **(c-f)**, the images used for MIP were subsets of those used in **(a)** or **(b)** to increase clarity. Image brightness and contrast for each panel are individually tuned for visualization. Scale bars: **(a)**, 1000 μm; **(b)**, 200 μm; **(c)**, **(d)**, **(e)**, **(f)**, 20 μm.

**Supplementary Figure 5.**
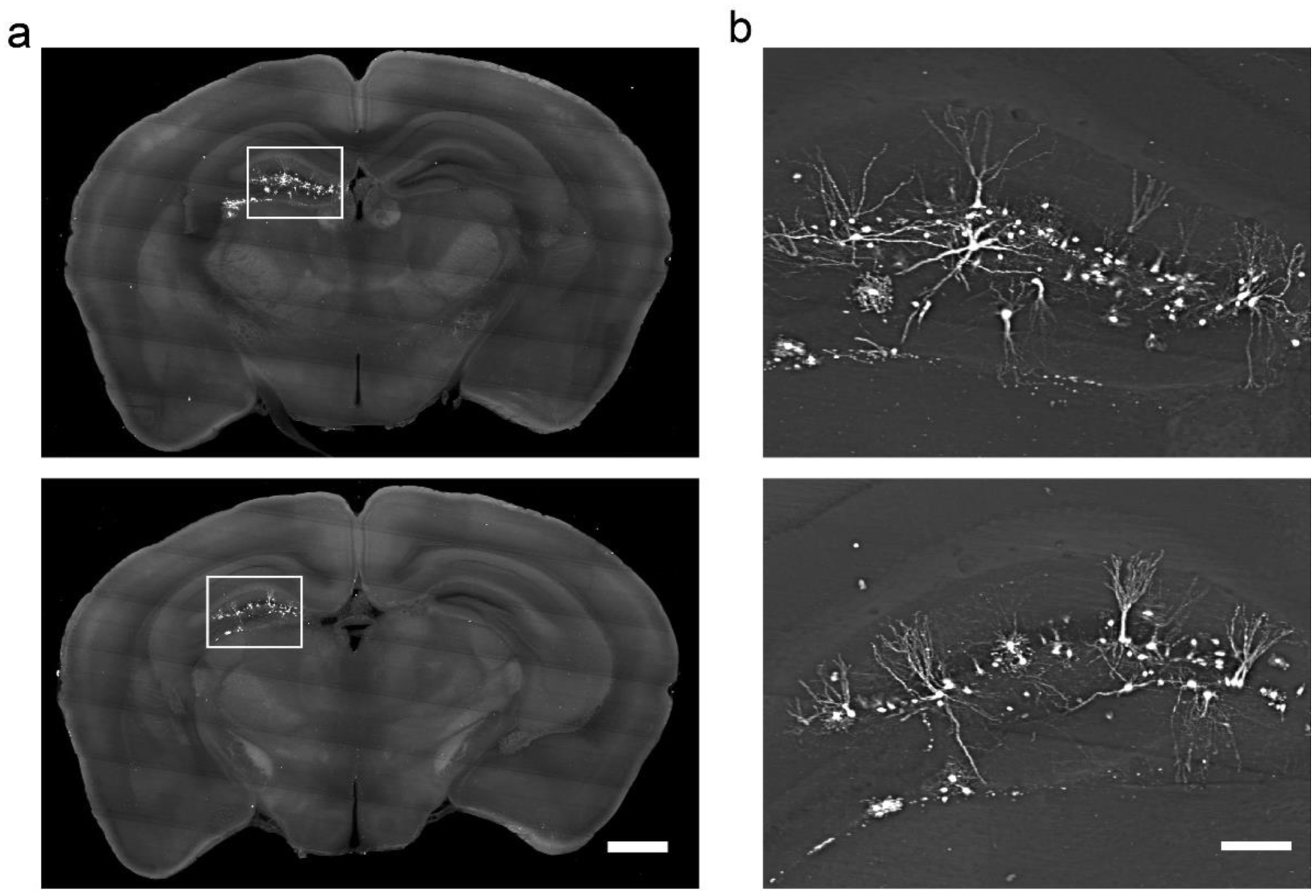
Hippocampal neurons sparsely labeled with Semliki Forest virus (SFV). **(a)** Brain sections showing sparsely labeled cells in the hippocampus 24 hours after SFV-EGFP injection. **(b)** Magnified view showing diverse neuronal morphology. Scale bars: **(a)**, 1 mm; **(b)**, 200 μm.

**Supplementary Figure 6.**
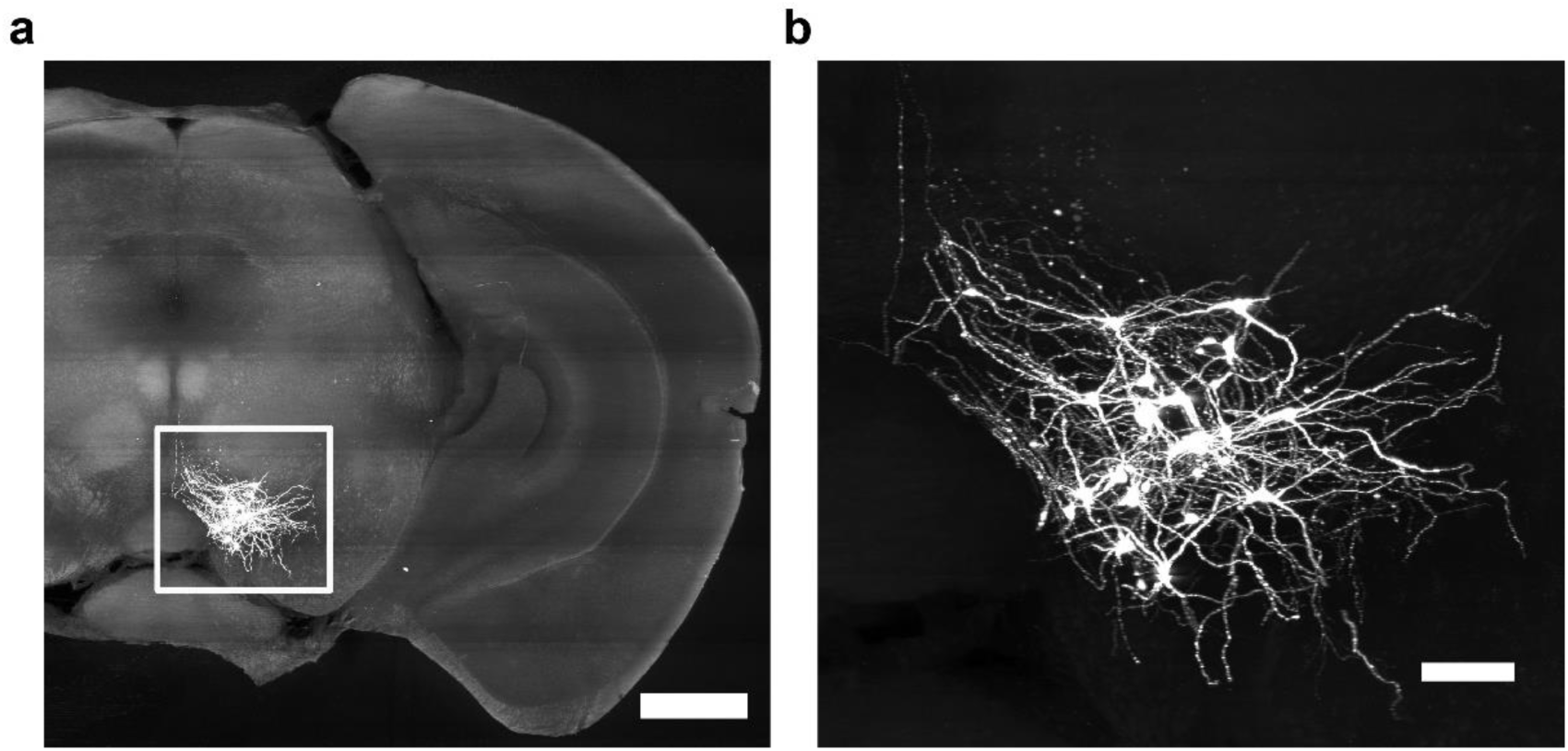
VTA dopaminergic neurons projecting to the nucleus accumbens revealed by cell-type specific tracing. **(a)** VTA dopaminergic neurons labeled by rVSV-EGFP-NR7A(A/RG) injected into the NAc in conjunction with rAAV-DIO-BFP-T2A-TVA injected into the VTA of a DAT-Cre mouse. **(b)** The morphology of EGFP-positive neurons clearly resolved by VISoR imaging. Scale bars: **(a)** 1 mm; **(b)** 200 μm.

**Supplementary Video 1**

VISoR image series from a 300-μm-thick Thy1-YFP brain slice captured at 100 fps, with smooth stage motion and synchronized scanning illumination. Fine structures of cortical neurons are revealed. Playback is at 0.1X real-time acquisition speed.

**Supplementary Video 2**

VISoR image series from a 300-μm-thick Thy1-YFP brain slice captured at 100 fps (top-left corner) with maximum intensity projections of these images after transformation and alignment. Playback is at the actual acquisition speed.

**Supplementary Video 3**

Navigation through an entire reconstructed Thy1-YFP mouse brain slice showing detailed neuronal structures.

**Supplementary Video 4**

Reconstructed whole brain of a Thy1-YFP mouse imaged by VISoR. The fluorescence intensity of each slice is adjusted according to its local contrast in order to enhance the global visualization clarity.

**Supplementary Video 5**

Dozens of dopaminergic neurons in the VTA traced from terminals in the NAc of a DAT-cre mouse, using a combination of VSV and AAV.

**Supplementary Video 6**

Dual-channel imaging shows that most CRH neurons (red) in the PVN are also c-fos positive (green) in the footshock-stimulated mouse but c-fos negative in the control mouse.

**Supplementary Video 7**

A large population of cells are activated (green, c-fos positive) in the medial amygdala (MeA) after footshock stimulation. Few of these activated cells are CRH expressing neurons (red).

## REFERENCES

1 Kleinfeld, D. et al. Large-scale automated histology in the pursuit of connectomes. The Journal of neuroscience: the official journal of the Society for Neuroscience 31, 16125–16138, doi:10.1523/JNEUROSCI.4077-11.2011 (2011).

2 Zingg, B. et al. Neural networks of the mouse neocortex. Cell 156, 1096–1111, doi:10.1016/j.cell.2014.02.023 (2014).

3 Oh, S. W. et al. A mesoscale connectome of the mouse brain. Nature 508, 207–214, doi:10.1038/nature13186 (2014).

4 Beier, K. T. et al. Circuit Architecture of VTA Dopamine Neurons Revealed by Systematic Input-Output Mapping. Cell 162, 622–634, doi:10.1016/j.cell.2015.07.015 (2015).

5 Zhang, S. et al. Selective attention. Long-range and local circuits for top-down modulation of visual cortex processing. Science 345, 660–665, doi:10.1126/science.1254126 (2014).

6 Wertz, A. et al. PRESYNAPTIC NETWORKS. Single-cell-initiated monosynaptic tracing reveals layer-specific cortical network modules. Science 349, 70–74, doi:10.1126/science.aab1687 (2015).

7 Xu, C. et al. Distinct Hippocampal Pathways Mediate Dissociable Roles of Context in Memory Retrieval. Cell 167, 961–972 e916, doi:10.1016/j.cell.2016.09.051 (2016).

8 Ye, L. et al. Wiring and Molecular Features of Prefrontal Ensembles Representing Distinct Experiences. Cell 165, 1776–1788, doi:http://dx.doi.org/10.1016/i.cell.2016.05.010 (2016).

9 Osten, P. & Margrie, T W. Mapping brain circuitry with a light microscope. Nature methods 10, 515–523, doi:10.1038/nmeth.2477 (2013).

10 Mayerich, D., Abbott, L. & McCormick, B. Knife-edge scanning microscopy for imaging and reconstruction of three-dimensional anatomical structures of the mouse brain. J Microsc 231, 134–143, doi:10.1111/j.1365-2818.2008.02024.x (2008).

11 Li, A. et al. Micro-optical sectioning tomography to obtain a high-resolution atlas of the mouse brain. Science 330, 1404–1408, doi:10.1126/science. 1191776 (2010).

12 Economo, M. N. et al. A platform for brain-wide imaging and reconstruction of individual neurons. Elife 5, e10566, doi:10.7554/eLife.10566 (2016).

13 Ragan, T et al. Serial two-photon tomography for automated ex vivo mouse brain imaging. Nature methods 9, 255–258, doi:10.1038/nmeth.1854 (2012).

14 Gong, H. et al. High-throughput dual-colour precision imaging for brain-wide connectome with cytoarchitectonic landmarks at the cellular level. Nature communications 7, doi:10.1038/ncomms12142 (2016).

15 Seiriki, K. et al. High-Speed and Scalable Whole-Brain Imaging in Rodents and Primates. Neuron 94, 1085–1100 e1086, doi:10.1016/j.neuron.2017.05.017 (2017).

16 Richardson, D. S., & Lichtman, J. W. Clarifying Tissue Clearing. Cell 162, 246–257, doi:10.1016/j.cell.2015.06.067 (2015).

17 Dodt, H. U. et al. Ultramicroscopy: three-dimensional visualization of neuronal networks in the whole mouse brain. Nature methods 4, 331–336, doi: 10.1038/nmeth1036 (2007).

18 Tomer, R., Ye, L., Hsueh, B. & Deisseroth, K. Advanced CLARITY for rapid and high-resolution imaging of intact tissues. Nature protocols 9, 1682–1697, doi:10.1038/nprot.2014.123 (2014).

19 Prevedel, R. et al. Simultaneous whole-animal 3D imaging of neuronal activity using light-field microscopy. Nature methods 11, 727–U161, doi:10.1038/Nmeth.2964 (2014).

20 Chung, K. et al. Structural and molecular interrogation of intact biological systems. Nature 497, 332–337, doi:10.1038/nature12107 (2013).

21 Renier, N. et al. Mapping of Brain Activity by Automated Volume Analysis of Immediate Early Genes. Cell 165, 1789–1802, doi:10.1016/j.cell.2016.05.007 (2016).

22 Wu, Y. et al. Spatially isotropic four-dimensional imaging with dual-view plane illumination microscopy. Nat Biotechnol 31, 1032–1038, doi:10.1038/nbt.2713 (2013).

23 Lemon, W. C. et al. Whole-central nervous system functional imaging in larval Drosophila. Nature communications 6, 7924, doi:10.1038/ncomms8924 (2015).

24 Wu, Y. et al. Inverted selective plane illumination microscopy (iSPIM) enables coupled cell identity lineaging and neurodevelopmental imaging in Caenorhabditis elegans. Proceedings of the National Academy of Sciences of the United States of America 108, 17708–17713, doi:10.1073/pnas. 1108494108 (2011).

25 Glaser, A. K. et al. Light-sheet microscopy for slide-free non-destructive pathology of large clinical specimens. Nature Biomedical Engineering 1, 0084, doi:10.1038/s41551-017-0084 https://www.nature.com/articles/s41551-017-0084#supplementary-information (2017).

26 Narasimhan, A., Umadevi Venkataraju, K., Mizrachi, J., Albeanu, D. F. & Osten, P. Oblique light-sheet tomography: fast and high resolution volumetric imaging of mouse brains. bioRxiv (2017).

27 Feng, G. et al. Imaging neuronal subsets in transgenic mice expressing multiple spectral variants of GFP. Neuron 28, 41–51 (2000).

28 Fuzesi, T., Daviu, N., Wamsteeker Cusulin, J. I., Bonin, R. P. & Bains, J. S. Hypothalamic CRH neurons orchestrate complex behaviours after stress. Nature communications 7, 11937, doi:10.1038/ncomms11937 (2016).

29 Lu, R. et al. Video-rate volumetric functional imaging of the brain at synaptic resolution. Nature neuroscience 20, 620–628, doi:10.1038/nn.4516 (2017).

30 Edelstein, A. D. et al. Advanced methods of microscope control using muManager software. J Biol Methods 1, doi:10.14440/jbm.2014.36 (2014).

31 Ehrengruber, M. U. et al. Recombinant Semliki Forest virus and Sindbis virus efficiently infect neurons in hippocampal slice cultures. Proceedings of the National Academy of Sciences of the United States of America 96, 7041–7046, doi:DOI 10.1073/pnas.96.12.7041 (1999).

32 Jia, F., Miao, H., Zhu, X. T. & Xu, F. Q. Pseudo-typed Semliki Forest virus delivers EGFP into neurons. J Neurovirol 23, 205–215, doi:10.1007/s13365-016-0486-8 (2017).

33 Beier, K. & Cepko, C. Viral Tracing of Genetically Defined Neural Circuitry. Jove-J Vis Exp, doi:UNSP e4253 10.3791/4253 (2012).

34 Chen, L. Y et al. Several residues within the N-terminal arm of vesicular stomatitis virus nucleoprotein play a critical role in protecting viral RNA from nuclease digestion. Virology 478, 9–17, doi:10.1016/j.virol.2015.01.021 (2015).

35 Backman, C. M. et al. Characterization of a mouse strain expressing Cre recombinase from the 3’ untranslated region of the dopamine transporter locus. Genesis 44, 383–390, doi:10.1002/dvg.20228 (2006).

36 Taniguchi, H. et al. A Resource of Cre Driver Lines for Genetic Targeting of GABAergic Neurons in Cerebral Cortex (vol 71, pg 995, 2011). Neuron 72, 1091–1091, doi:10.1016/j.neuron.2011.12.010 (2011).

37 Madisen, L. et al. A robust and high-throughput Cre reporting and characterization system for the whole mouse brain. Nature neuroscience 13, 133–U311, doi:10.1038/nn.2467 (2010).

38 Jia, F., Zhu, X. T. & Xu, F. Q. A single adaptive point mutation in Japanese encephalitis virus capsid is sufficient to render the virus as a stable vector for gene delivery. Virology 490, 109–118, doi:10.1016/j.virol.2016.01.001 (2016).

39 Yang, B. et al. Single-Cell Phenotyping within Transparent Intact Tissue through Whole-Body Clearing. Cell 158, 945–958, doi:10.1016/j.cell.2014.07.017 (2014).

40 Schindelin, J., Rueden, C. T., Hiner, M. C. & Eliceiri, K. W. The ImageJ ecosystem: An open platform for biomedical image analysis. Mol Reprod Dev 82, 518–529, doi:10.1002/mrd.22489 (2015).

41 Liu, C., Yuen, J. & Torralba, A. SIFT Flow: Dense Correspondence across Scenes and Its Applications. Ieee T Pattern Anal 33, 978–994, doi:10.1109/Tpami.2010.147 (2011).

42 Schaefer, S., McPhail, T. & Warren, J. Image deformation using moving least squares. Acm T Graphic 25, 533–540, doi:Doi10.1145/1141911.1141920 (2006).

43 Durand, F. & Dorsey, J. Fast bilateral filtering for the display of high-dynamic-range images. Acm T Graphic 21, 257–266 (2002).

